# starmap: Immersive visualisation of single cell data using smartphone-enabled virtual reality

**DOI:** 10.1101/324855

**Authors:** Andrian Yang, Yu Yao, Jianfu Li, Joshua W. K. Ho

**Affiliations:** Victor Chang Cardiac Research Institute, Darlinghurst, NSW 2010, Australia; University of New South Wales, Sydney, Australia

## Abstract

We report a new smartphone-enabled virtual reality (VR) program, starmap (https://vccri.github.io/starmap/), that enables immersive visualisation of single-cell data for hundreds of thousands of cells using a mobile-enabled web browser and low-cost VR head mount device.

---

Advances in single-cell RNA-seq technology^1,2^, flow cytometry^3^ and mass cytometry^4^ has enabled the expression profiling of a large number of genes and proteins for hundreds of thousands of individual cells. This has opened up opportunities to explore cellular heterogeneity and has application in almost all disciplines of biology and medicine. Dimensionality reduction methods, such as Principal Component Analysis (PCA) or other non-linear methods^5^, can reduce the large number of features into lower dimensions to allow for efficient analysis of data. The data can then be visualised into in two or three dimensions on a computer screen or on a printed page. However, displaying and understanding of thousands of points are often challenging enough, let alone hundreds of thousands of points in the case of single cell data.

An effective visualisation should allow a viewer to not only assess the high-level clustering structure of cells, but also allow viewers to seamlessly ‘zoom’ into the data to discover fine local grouping of single cells, and to explore their gene/protein expression profiles. This type of visualisation is not easy to achieve using standard computer flat-screen visualisation.

starmap breaks new ground on large scale data visualisation in two ways. First, it introduces a scalable visual design that combines the benefit of a three-dimensional (3D) scatter plot (Figure 1a) for exploring clustering structure and the benefit of star plots^6^ (also known as radar chart) for multivariate visualisation of an individual cell (Figure 1b). Second, starmap is designed to utilise low-cost virtual reality (VR) headsets, such as Google Cardboard and Daydream, to enable widespread adoption of VR data visualisation. We reason that an immersive visual experience will likely improve the navigation and exploration of hundreds of thousands of cells.

**Figure 1.**
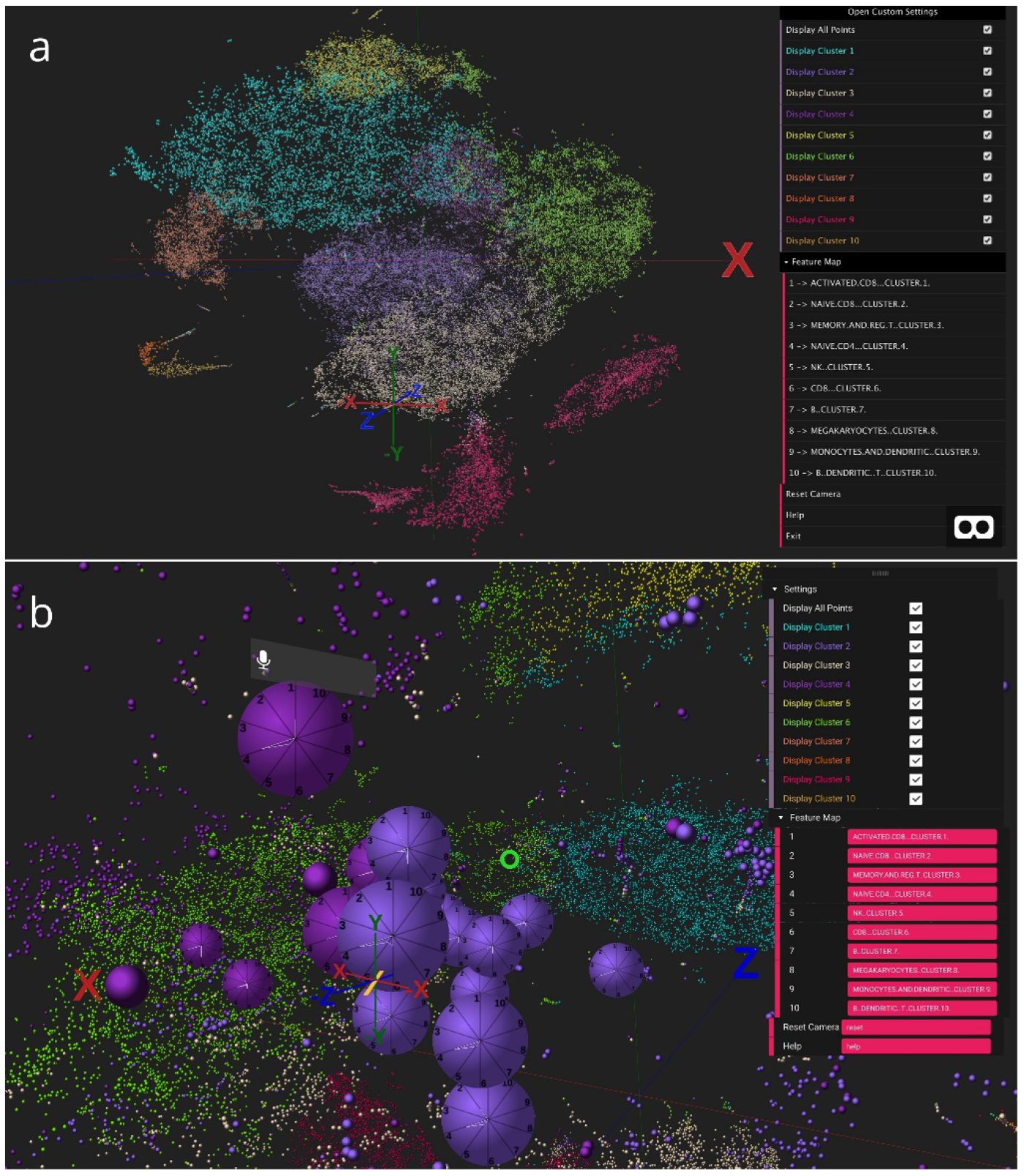
starmap visualisation of a single-cell RNA-seq data containing 68,000 peripheral blood mononuclear cells (**a**) Overview of the 3D scatter plot with visualised in non-VR mode (**b**) Star plot visualisation showing the relative expression of gene modules for each cell in VR mode.

starmap, is designed to visualise single cell data primarily through a web browser on either a computer or a smartphone. starmap is built using web frameworks designed for creation of 3D and VR experiences, such as A-Frame and Three.js, which are cross-platform and can be adapted to computer screens and VR devices. Input data for starmap, in the form the 3D coordinates, cluster assignment, and value of up to 12 features per cell in a comma-separated values format, can be uploaded from a local file on your device or from third-party cloud storage, such as Google Drive, Microsoft OneDrive or iCloud Drive, depending on the support of the mobile smartphone’s operating system. starmap offers a number of options for navigation through the VR space. Users are able to use a wireless keyboard or wireless hand-held remote controller in order to scale, rotate and navigate within the VR space. starmap also offers a voice control feature that allows user to start and end custom animation for navigating the VR space

For the star plot visualisation, starmap supports up to 12 radial coordinates which emanates from the centre to the circumference of the circle, with the coordinate indicating the expression level of a particular feature. The feature typically represents biological feature such a gene, in the case of RNA-seq data, or a protein, in the case of flow cytometry data. However, the feature can also represent can also be a loading value for a principal component, the mean expression of a group of genes or proteins, or some other aggregate representation of related features.

The starmap website contains two example data sets based on previously published single-cell RNA-seq data^7^ and flow cytometry data^8^ (see **Supplementary Methods** for details of data processing). The source code and sample data are available at https://github.com/VCCRI/starmap.

## Author contributions

A.Y. and J.W.K.H. initiated the idea, designed the prototype, and wrote the paper. Y.Y. and J.L. implemented and tested starmap. All authors revised and approved the final version of the manuscript.

